# Phantom histories of misspecified pasts

**DOI:** 10.1101/2020.06.26.173963

**Authors:** Alexander Platt, Daniel N. Harris

## Abstract

The observation that even a tiny sample of genome sequences from a natural population contains a plethora of information about the history of the population has enticed researchers to use these data to fit complex demographic histories and make detailed inference about the changes a population has experienced through time. Unfortunately, the standard assumptions required to make these inferences are often violated by natural populations in such ways as to produce specious results. This paper examines two phenomena of particular concern: when a sample is drawn from a single sub-population of a larger meta-population these models infer a spurious recent population decline, and when a genome contains loci under weak or recessive purifying selection these models infer a spurious recent population expansion.

## Introduction

Coalescent theory gives us a powerful tool linking observable patterns of genetic diversity in a sample of individuals to historical properties of the populations from which they descend (*e*.*g*. [1, 2]). These approaches have allowed us to estimate important phenomena such as population sizes, migration rates among populations, and historical changes in these governing parameters. Recent methods implementing Hidden Markov Models to leverage genome sequencing data (*e*.*g*. [3, 4, 5, 6]) and the lengths of shared segments between individuals (*e*.*g*. [7, 8]) have extended the capabilities of these analyses to fit models with dozens of parameters representing fine-scale changes in populations’ parameters through time.

Problems, though, can arise, in interpreting the results when analytic models do not account for some of the important forces contributing to the composition of a population sample. These do not end at a constant scalar relating effective and consensus population sizes, nor manifestations of brief periods of deviation in population structure [3] or natural selection [9] as corresponding periods of deviation in population size. Other forms of model misspecification, can produce more fundamental difficulties where evidence of dramatic population expansion or contraction arise from populations at complete equilibrium and is not associated with *any* change experienced by the population. This paper examines two such conditions. In the first, the mere existence of other, un-sampled populations is sufficient to generate a signature of recent population contraction even if none of the populations have ever changed size and all of the parameters describing migration among them have always remained fixed [10]. The second is ongoing purifying selection with mutations that are either recessive or weakly deleterious. This generates a signature of population expansion, even without anything changing regarding the mutational spectrum, fitness landscape, or underlying genetic architecture.

## Results

### Un-sampled populations

We simulated 100 replicate samples of two chromosomes taken from the central population of a nine-population stepping-stone model [11] where each population was comprised of 10,000 haploid individuals and where each population exchanged a fixed percentage of its individuals with the adjacent populations every generation. These were entirely neutral simulations of populations with fixed demographic structure. None of the parameters changed at any point during a simulation. When migration between populations is on the order of a single migrant entering a population each generation (figure 1) the simulations show a consistent signature of population contraction with recent population size estimates one tenth the size of older estimates. With decreasing migration rates the magnitude of the signature of population contraction grows and its onset is pushed farther back in time (supplemental figures S1-S3). As expected [12], much higher migration rates cause the entire meta-population to behave as a single, large, panmictic population with no signature of population contraction (supplemental figure S4). With no migration at all the sampled population behaves as a single, small, panmictic population and again exhibits no signature of population contraction (supplemental figure S5).

**Figure 1:**
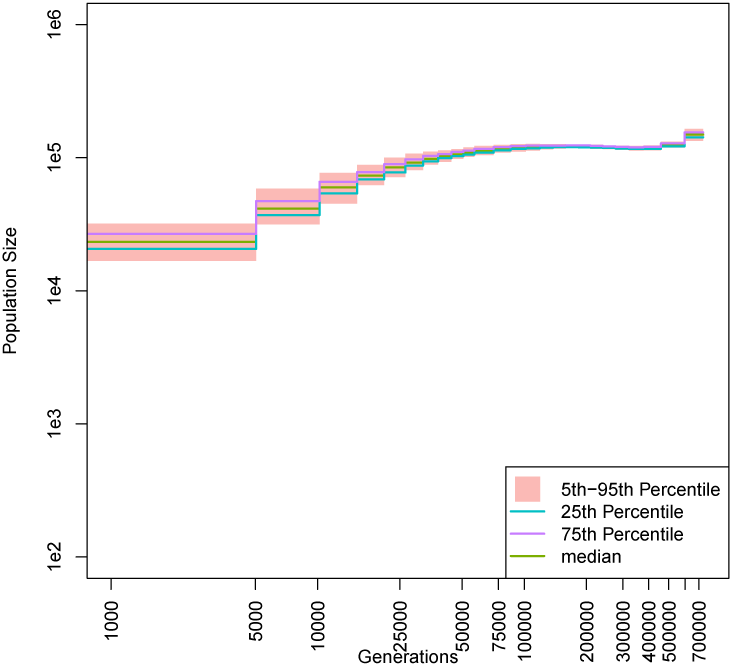
Inferred historic population size in presence of un-sampled populations. Estimates of effective population sizes from a sample of two chromosomes drawn from the central of nine linearly arrayed populations. Each population has a fixed size of 10^4^ and the per-generation migration rate between adjacent populations is a constant 10^−4^.

### Weak or recessive purifying selection

We simulated sets of 100 replicate constant-sized Wright-Fisher populations of 3,000 diploid individuals experiencing weak purifying selection. With half of all mutations being neutral and half conferring an additive 0.1% reduction in fitness we find a signature of population expansion with recent population size estimates an order of magnitude larger than older ones (figure 2). These mutations are weak enough that they are approaching the lower limit of what natural selection can act on [13]. Mutations that are more deleterious and systems where a higher proportion of mutations are deleterious both give rise to even starker signatures of population expansion (supplemental figures S6-S10).

**Figure 2:**
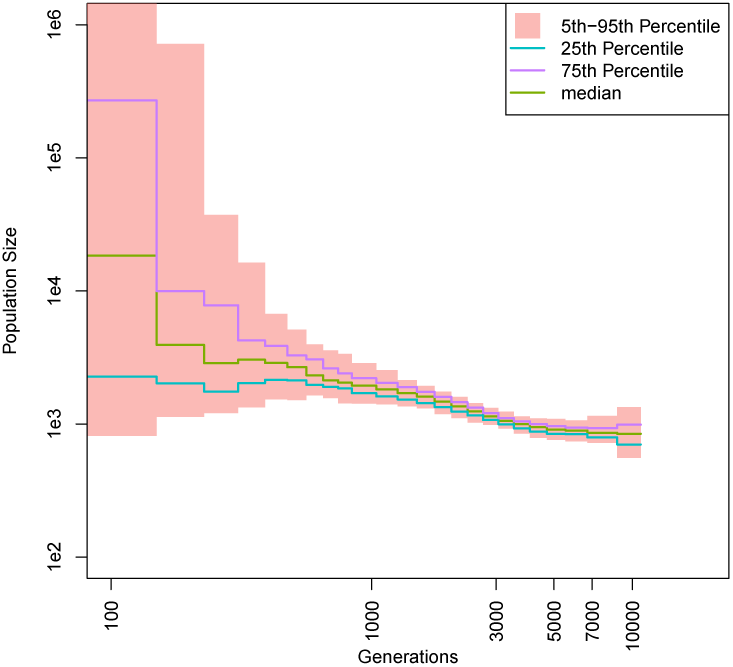
Inferred historic population size in presence of mildly deleterious mutations. Estimates of effective population sizes from a sample of two chromosomes drawn from a population of 3,000 individuals experiencing weak purifying selection. Each new mutation has a 50% chance of conferring a 0.1% reduction of fitness.

We also simulated sets of 100 replicate constant-sized Wright-Fisher populations of 3,000 diploid individuals where in addition to neutral mutations, a fraction of new mutations were lethal when homozygous but neutral when heterozygous. These simulations produced signatures of recent population expansion as well (figure 3). The effect is still present, though less consistently, when as few as 10% of mutations are recessive lethal (supplemental figure S11).

**Figure 3:**
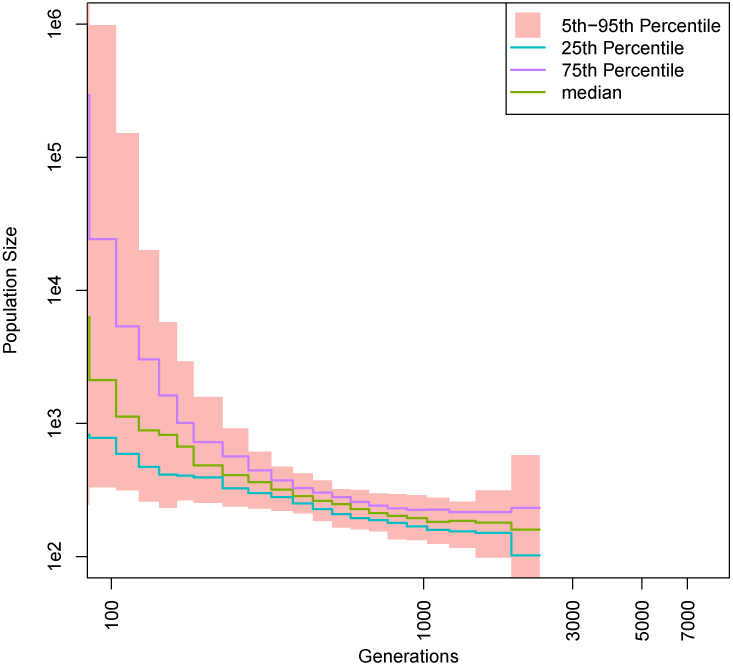
Inferred historic population size in presence of recessive lethal mutations. Estimates of effective population sizes from a sample of two chromosomes drawn from a population of 3,000 individuals where each new mutation has a 50% chance of incurring lethality when homozygous.

## Conclusions

These phantom signatures of historic changes in population size are not the result of any sort of implementation or computational difficulty. Rather, they stem from fundamental model misspecifications. As discussed recently in [14], the concept of an effective population size (*N*_*e*_) has been a powerful one that has been defined and interpreted in many different ways throughout the history of population genetics. One particular definition of *N*_*e*_, the coalescent effective population size 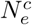, is the linear scaling of the Kingman coalescent [15] that yields the expected distribution of times of shared ancestry among individuals within the sample [16]. This is the concept of population size that is being estimated from coalescent-based demographic inference algorithms. Deviations from a constantly-scaled Kingman coalescent history of a sample are being interpreted as changes in historical population size. Difficulty arises from conditions for which a sample’s genealogy is not expected to converge to a Kingman coalescent. If there is no single re-scaling that recovers a Kingman coalescent, algorithms are forced to introduce temporally-shifting population sizes to explain the inferred distribution of coalescent events. This is why even with completely stationary phenomena such as the presence of un-sampled populations or mildly deleterious (or recessive) mutations, coalescent-based inference of demographic history produces artifacts of dynamic changes in effective population size.

Intuitively, for the example of un-sampled populations, one can imagine that individuals who share recent common ancestors do so by a process dominated by the local parameters of the population in which they were sampled. If individuals do not share a recent common ancestor, however, the probability increases that they have ancestors who belonged to different populations, and the process that describes their relationship to each other is one that involves multiple populations’ worth of individuals as well as the rate of migration among them. This time-varying signature from a static demographic structure is a distinct phenomenon from the transient increases in 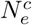 due to transient increases in population structure discussed in [3], but is similar to the effect shown in [10] for symmetrical island models.

With weak or recessive deleterious mutations, new mutations are not immediately removed from the population. While they segregate, carriers of these alleles are free to contribute to the ancestry pool of subsequent generations. Over time, however, natural selection acts to remove these variants, reducing the number of potential ancestors over longer time periods. In both cases, at any and every point in time, the process describing the recent history of a sample is distinct from the process describing its more distant past, though neither of these processes involve any change in the population at any point in time. “Recent” and “distant” history are forever relative to the time of sampling as the genealogy will never have the simple form of a Kingman coalescent.

This problem is then compounded when comparing samples taken at different time points, such as a modern sample and ancient DNA. In any of the above examples, if the ancient sample is drawn from a population directly ancestral to the modern sample the two will share identical patterns of expansion or contraction, but do so relative to their distinct sample times producing diverged historical effective population sizes. If their shared history includes periods of true demographic variation (such as a population bottleneck or selective sweep), those signals would appear on an absolute timescale. No translation or scaling of the inferred historical 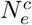 would align the two trajectories, even though they represent the same population.

Un-sampled populations, mildly deleterious mutations, and recessive deleterious mutations are all examples of phenomena that cause a population to deviate from the Kingman coalescent, and yet are wide-spread throughout the natural world. Entire species rarely behave as a single panmictic population and it is not reasonable to expect that any researcher will sample all of a species’ populations. Organisms such as *Arabidopsis thaliana* with reduced non-coding portions of the genome [17] and population structure that can be characterized primarily by isolation by distance [18] are likely to be particularly recalcitrant to accurate inference and may comprise a worst-case scenario.

Going forward, coalescent-based algorithms for demography inference will require more care and attention to implementation and interpretation. Modern methods allow us to accurately estimate historical coalescent intensities. Simply summarizing those intensities as a function of a time-varying effective population size, however, can lead to greater confusion than insight. Considering the undesired confounding from natural selection, whenever possible researchers should start by masking genomes to enrich for regions that are far removed from regions known to experience selective constraint. Considering population structure, it will be necessary to develop new analytical methods flexible enough to directly account for demographic histories that are far more complex than the current state of the art. Until that point, results from these kinds of inference must be treated skeptically, as the values represented will reflect a delicate balance of multiple different forces occurring across different time periods and likely varying in composition and strength across populations.

## Methods

Each simulation used constant mutation and recombination rates of 10^−8^ per-base, per-generation. We used coalescent simulation (as implemented in msprime version 0.7.3 [19]) to create pairs of 50Mb chromosomes from each replicate for the neutral stepping stone scenarios. We used Wright-Fisher simulations (as implemented in `SLiM3.3.1` [20]) for scenarios involving natural selection. The Wright-Fisher simulations were initialized from neutral expectations and run forward in time for 30,000 (10*N*) generations before sampling two 100 Mb chromosomes from each. Mutations in `SLiM` were sampled from a fixed fitness effect distribution, and all mutations that became fixed in the population were not removed from the simulation by setting “convertToSubstitution = F”. All simulated variants with a physical position that appears more than once in a single simulation were removed from the analysis. We used `MSMC` [5] version with default settings to infer the demographic history of each sample. Each `MSMC` input file was formatted using `generate_multihetsep`.py from `msmc-tools`. MSMC output 40 time steps, but different time intervals for each simulation replicate. Therefore, we used the median time value at each time interval when forming the step plots of population size estimates.

## Acknowledgements

This work was supported by the National Institutes of Health (R35 GM134957-01 to A.P. and 5T32DK007314-39 to D.N.H) and the American and the Alcohol Education Research Council Diabetes Association Pathway to Stop Diabetes (1-19-VSN-02 to A.P.). We would like to thank Iain Mathieson and Sara Mathieson for valuable feedback.

## Supplementary Material

**Figure S1:**
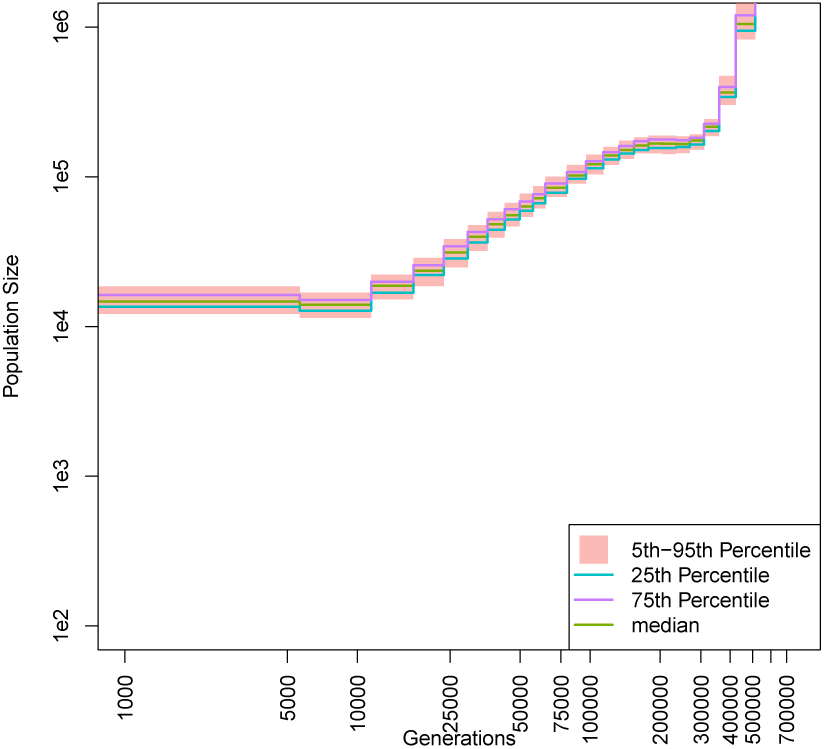
Inferred historic population size in presence of un-sampled populations. Estimates of effective population sizes from a sample of two chromosomes drawn from the central of nine linearly arrayed populations. Each population has a fixed size of 10^4^ and the per-generation migration rate between adjacent populations is a constant 10^−5^.

**Figure S2:**
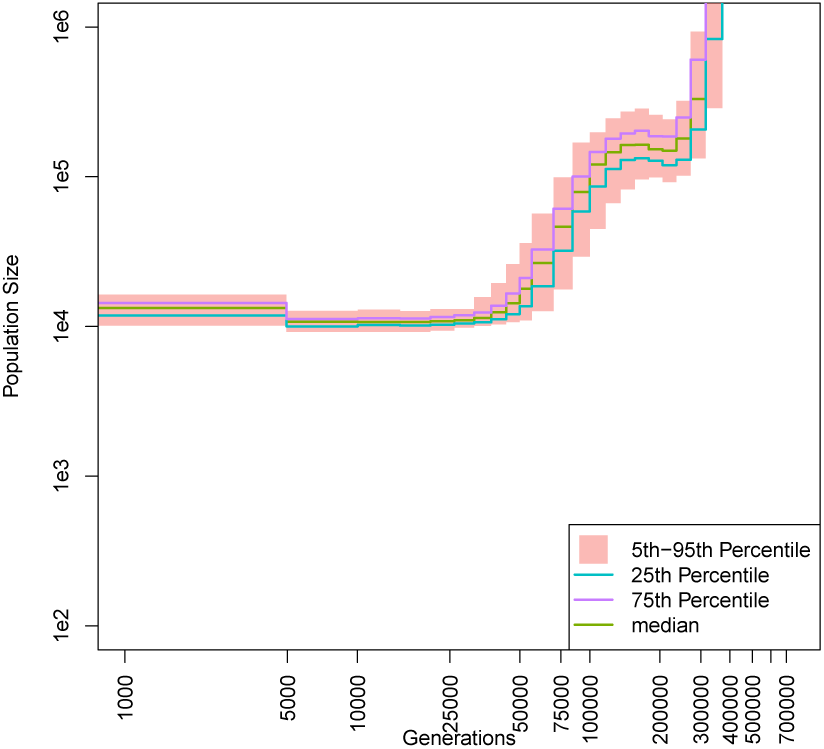
Inferred historic population size in presence of un-sampled populations. Estimates of effective population sizes from a sample of two chromosomes drawn from the central of nine linearly arrayed populations. Each population has a fixed size of 10^4^ and the per-generation migration rate between adjacent populations is a constant 10^−6^.

**Figure S3:**
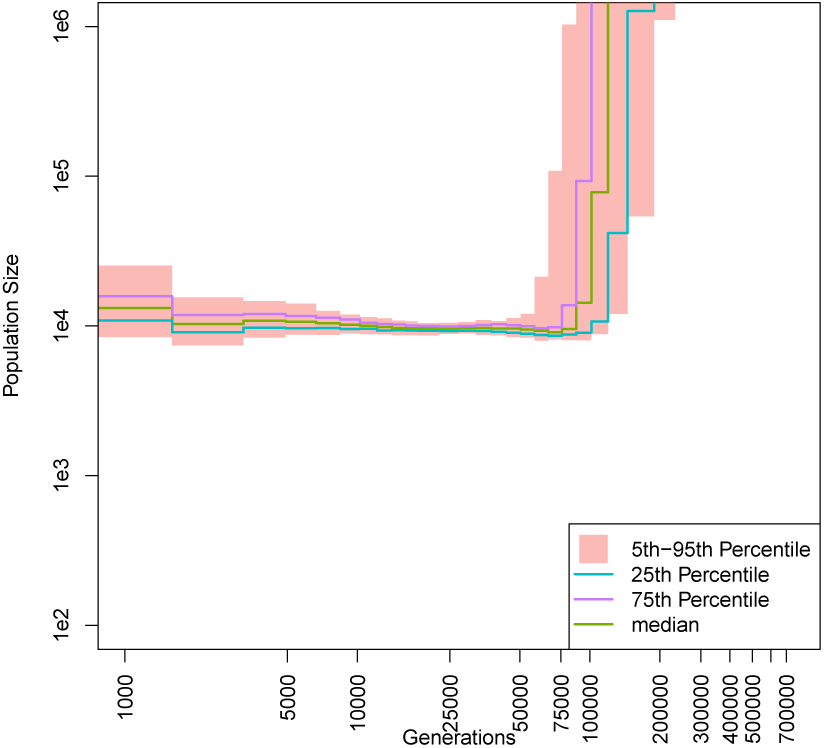
Inferred historic population size in presence of un-sampled populations. Estimates of effective population sizes from a sample of two chromosomes drawn from the central of nine linearly arrayed populations. Each population has a fixed size of 10^4^ and the per-generation migration rate between adjacent populations is a constant 10^−7^.

**Figure S4:**
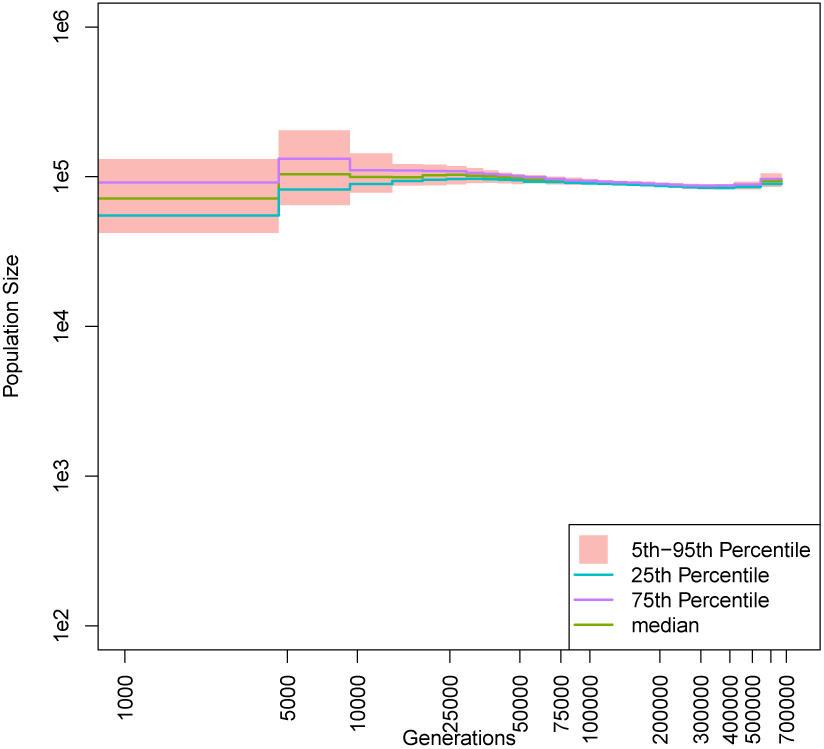
Inferred historic population size in presence of un-sampled populations. Estimates of effective population sizes from a sample of two chromosomes drawn from the central of nine linearly arrayed populations. Each population has a fixed size of 10^4^ and the per-generation migration rate between adjacent populations is a constant 10^−3^.

**Figure S5:**
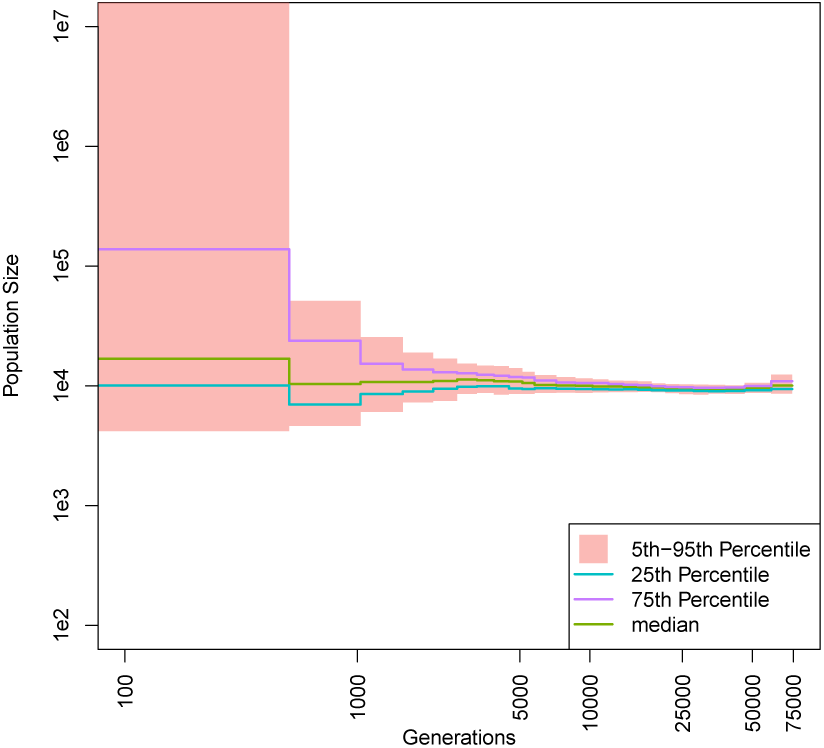
Inferred historic population size in presence of un-sampled populations. Estimates of effective population sizes from a sample of two chromosomes drawn from the central of nine linearly arrayed populations. Each population has a fixed size of 10^4^. There is no migration among populations.

**Figure S6:**
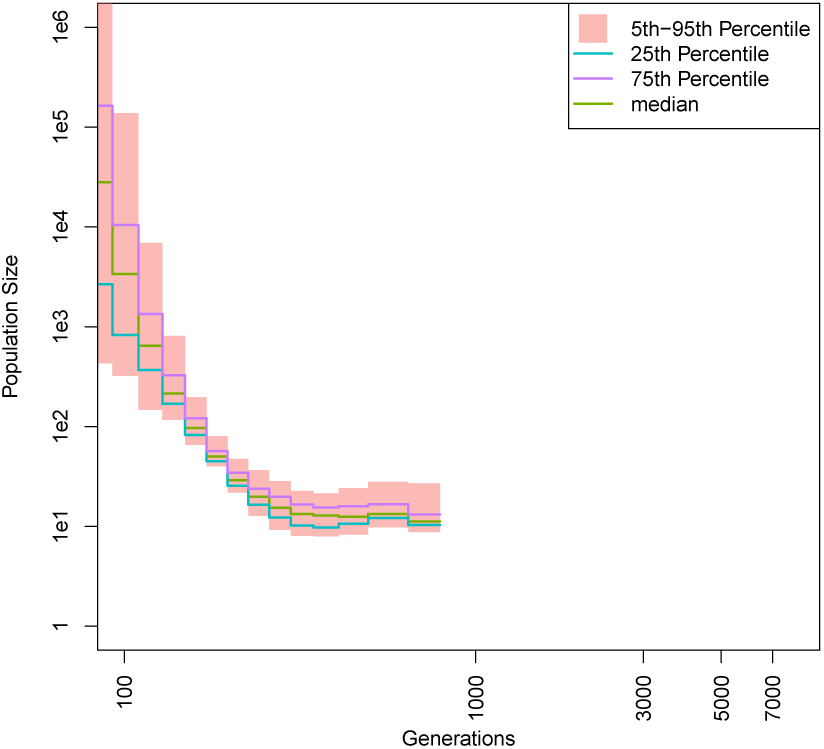
Inferred historic population size in presence of deleterious mutations. Estimates of effective population sizes from a sample of two chromosomes drawn from a population of 3,000 individuals experiencing purifying selection. Each new mutation confers a 1% reduction of fitness.

**Figure S7:**
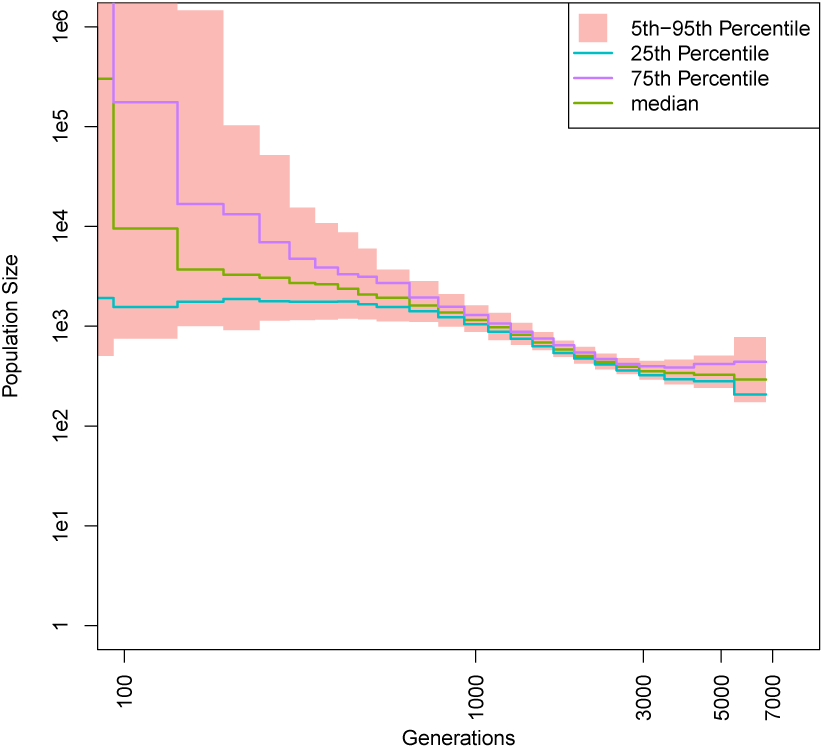
Inferred historic population size in presence of deleterious mutations. Estimates of effective population sizes from a sample of two chromosomes drawn from a population of 3,000 individuals experiencing purifying selection. Each new mutation confers a 0.1% reduction of fitness.

**Figure S8:**
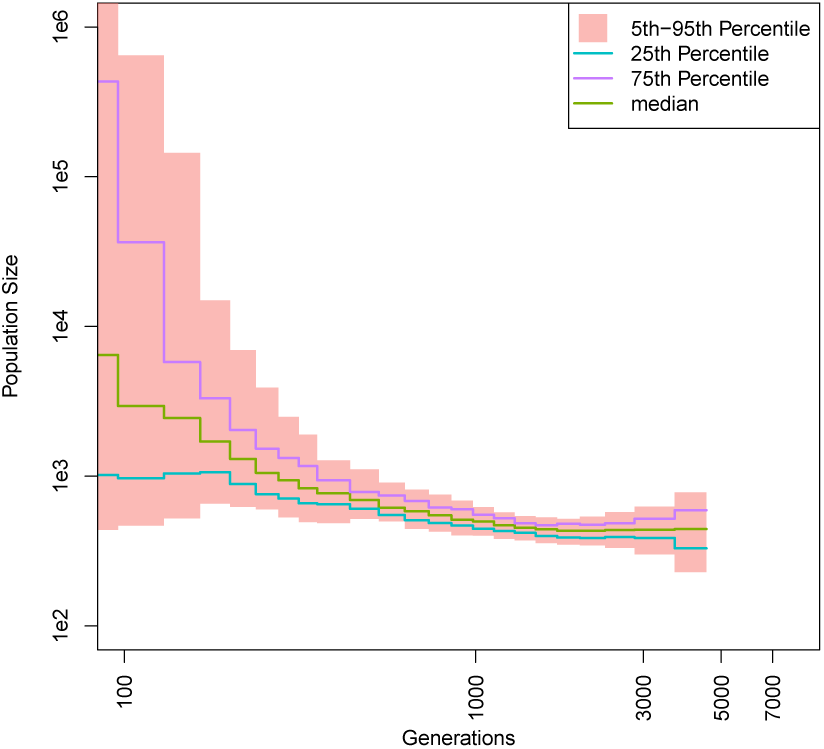
Inferred historic population size in presence of deleterious mutations. Estimates of effective population sizes from a sample of two chromosomes drawn from a population of 3,000 individuals experiencing purifying selection. Each new mutation has a 50% chance of conferring a 1% reduction of fitness.

**Figure S9:**
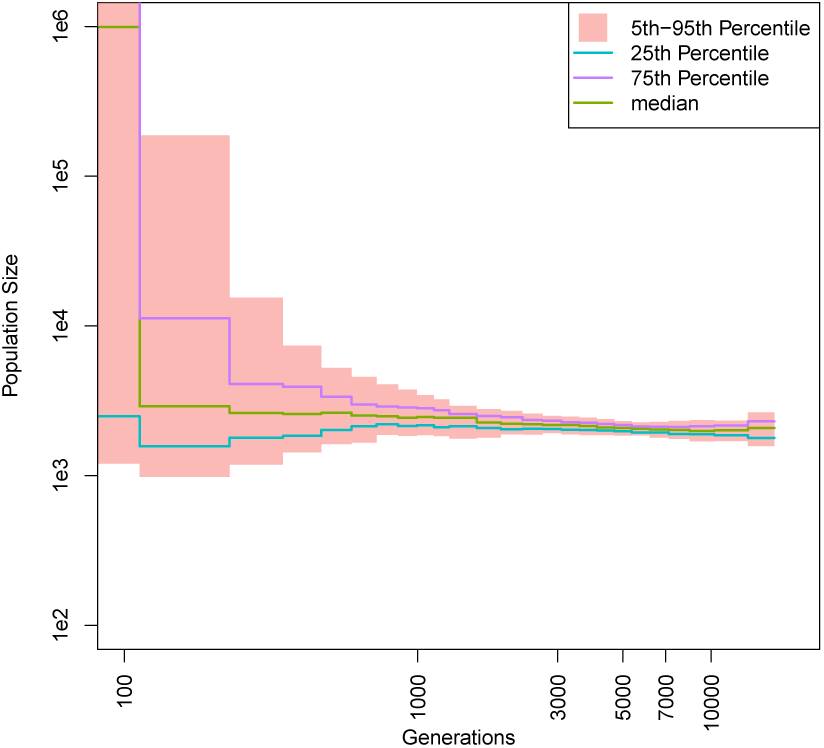
Inferred historic population size in presence of deleterious mutations. Estimates of effective population sizes from a sample of two chromosomes drawn from a population of 3,000 individuals experiencing purifying selection. Each new mutation has a 10% chance of conferring a 1% reduction of fitness.

**Figure S10:**
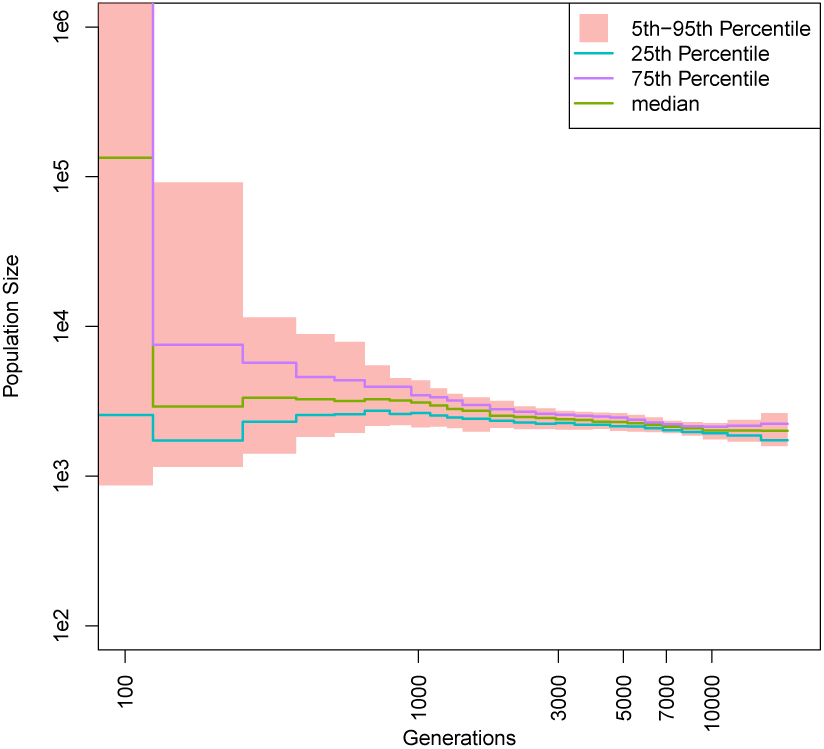
Inferred historic population size in presence of mildly deleterious mutations. Estimates of effective population sizes from a sample of two chromosomes drawn from a population of 3,000 individuals experiencing weak purifying selection. Each new mutation has a 10% chance of conferring a 0.1% reduction of fitness.

**Figure S11:**
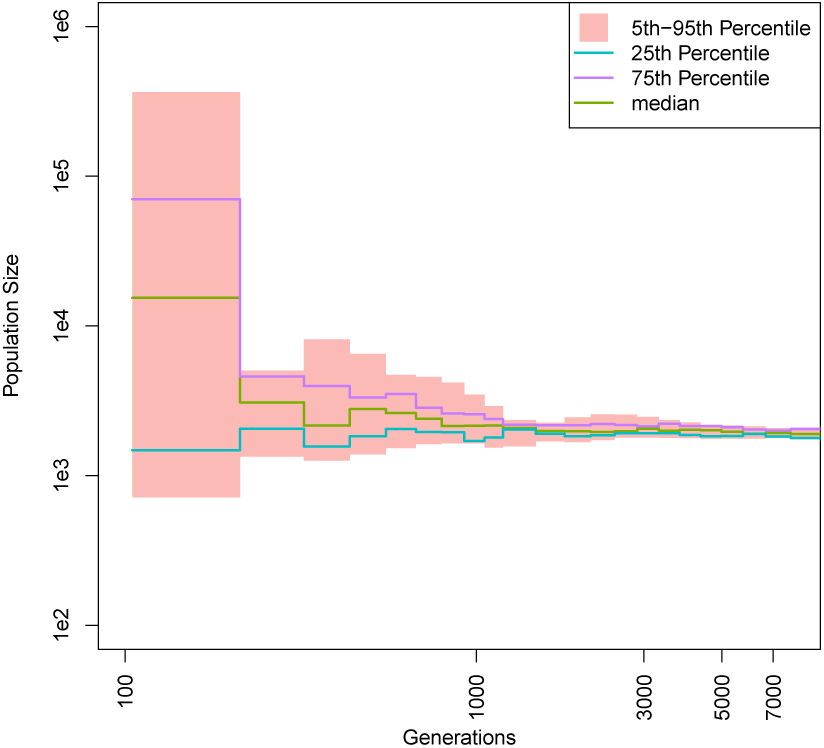
Inferred historic population size in presence of recessive lethal mutations. Estimates of effective population sizes from a sample of two chromosomes drawn from a population of 3,000 individuals where each new mutation has a 10% chance of incurring lethality when homozygous.

